# Common spatiotemporal processing of visual features shapes object representation

**DOI:** 10.1101/459214

**Authors:** Paolo Papale, Monica Betta, Giacomo Handjaras, Giulia Malfatti, Luca Cecchetti, Alessandra Rampinini, Pietro Pietrini, Emiliano Ricciardi, Luca Turella, Andrea Leo

## Abstract

Biological vision relies on representations of the physical world at different levels of complexity. Relevant features span from simple low-level properties, as contrast and spatial frequencies, to object-based attributes, as shape and category. However, how these features are integrated into coherent percepts is still debated. Moreover, these dimensions often share common biases: for instance, stimuli from the same category (e.g., tools) may have similar shapes. Here, using magnetoencephalography, we revealed the temporal dynamics of feature processing in human subjects attending to pictures of items pertaining to different semantic categories. By employing Relative Weights Analysis, we mitigated collinearity between model-based descriptions of stimuli and showed that low-level properties (contrast and spatial frequencies), shape (medial-axis) and category are represented within the same spatial locations early in time: 100-150ms after stimulus onset. This fast and overlapping processing may result from independent parallel computations, with categorical representation emerging later than the onset of low-level feature processing, yet before shape coding. Categorical information is represented both before and after shape also suggesting a role for this feature in the refinement of categorical matching.

## Introduction

To make sense of the surrounding environment, our visual system relies on different transformations of the retinal input^1^. Just consider Figure 1A. As any natural scene, this image is defined by a specific content of edges and lines. However, biological vision evolved to disclose the layout of discrete objects, hence the two giraffes in the foreground emerge as salient against the background, and the distinct contents pertaining to edges, shape, texture, and category contribute together to object perception.

**Figure 1.**
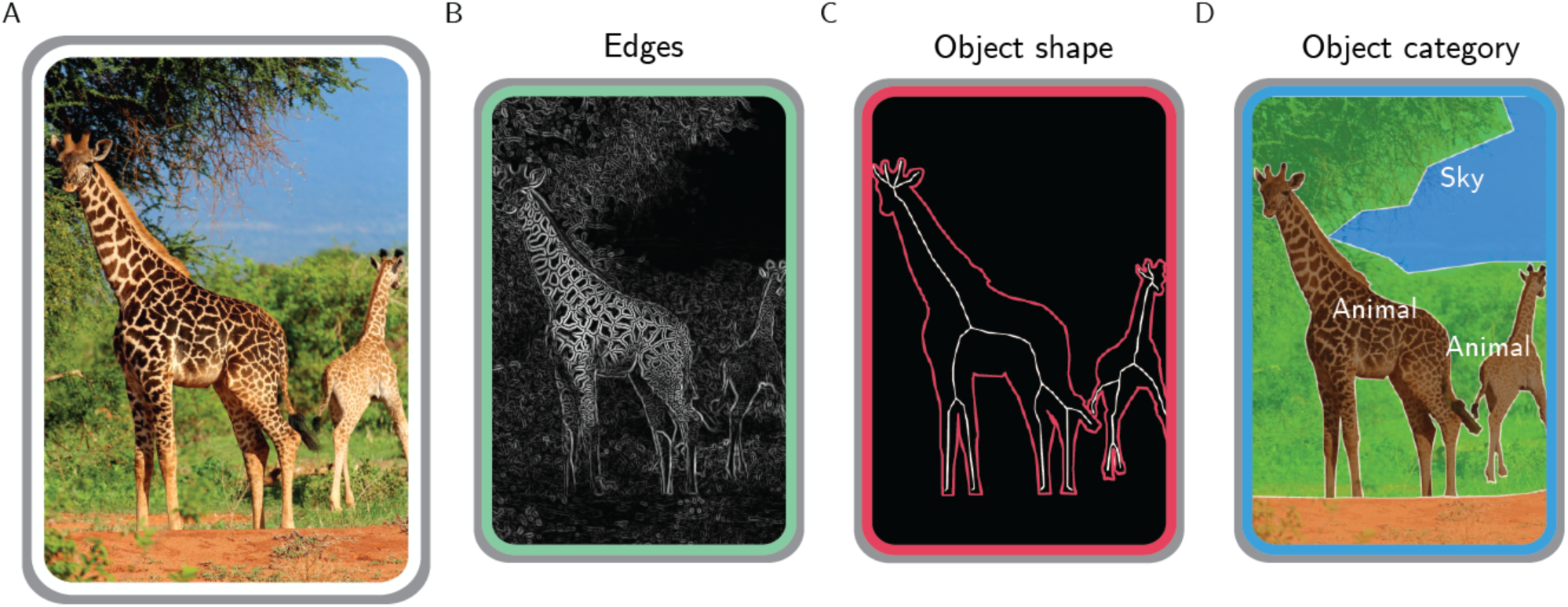
Different representations of a natural image. A real-world scene (A), depicting two giraffes in the savannah, can be defined by its edges (B), by the shape of the giraffes (C) and also by the categorical information it conveys (D). Photo taken from http://pixabay.com, released under Creative Commons CC0 license.

Actually, each feature of Figures 1B-D is processed across the whole visual system. The primary visual cortex provides an optimal encoding of natural image statistics based on local contrast,orientation and spatial frequencies ^2,3^, and these low-level features significantly correlate with brain activity in higher-level visual areas^4,5^. Nonetheless, occipital, temporal and parietal modules also process object shape^6-8^ and categorical knowledge^9-11^.

Although all these features are relevant to our brain, their relative contribution in producing discrete and coherent percepts has not yet been clarified. In general, these different dimensions are interrelated and share common biases (i.e., are collinear), thus limiting the capability to disentangle their specific role^12^. For instance, categorical discriminations can be driven either by object shape (e.g., tools have peculiar outlines) or spatial frequencies (e.g., faces and places have specific spectral signatures: ^13^). Consequently, object shape and category are processed by the same regions across the visual cortex, even when using a balanced set of stimuli^14^. Even so, the combination of multiple feature-based models appears to describe underlying object representations at a neural level better than the same models tested in isolation. For instance, a magnetoencephalography (MEG) study found that combining low-level and semantic features improves the prediction accuracy of brain responses to viewed objects, thus suggesting that semantic information integrates with visual features during the temporal formation of object representations^15^.

Here, we combined model-based descriptions of pictures, MEG brain activity patterns and a statistical procedure (Relative Weights Analysis; RWA, ^16^) that mitigate the effects of common biases across different dimensions, to investigate the spatiotemporal dynamics of object processing and ultimately determine the relative contribution across space and time of multiple feature-based representations – i.e., low-level, shape and categorical features - in producing the structure of what we perceive. First, the low-level description of the stimuli was grounded on features extracted by the early visual cortex (i.e., image contrast and spatial frequencies). Second, since shape is critical to interact with the surrounding environment^17^, we relied on a well-assessed, physiologically-motivated description of shape, i.e., the medial axis ^18^. Finally, objects were also distinctively represented according to their superordinate categories.

To anticipate, we observed fast (100-150ms) and overlapping representations of low-level properties (contrast and spatial frequencies), shape (medial-axis) and category in posterior sensors. These results may be interpreted as macroscale dynamics resulting in independent parallel processing, and may also suggest a role for shape in the refinement of categorical matching.

## Results

We employed the Relative Weights Analysis^16^ to reveal the proportional contribution of low-level, shape and category feature models in predicting time resolved representational geometries derived from MEG data, recorded from subjects attending to pictures representing thirty different stimuli from six semantic categories (Figure 2).

**Figure 2.**
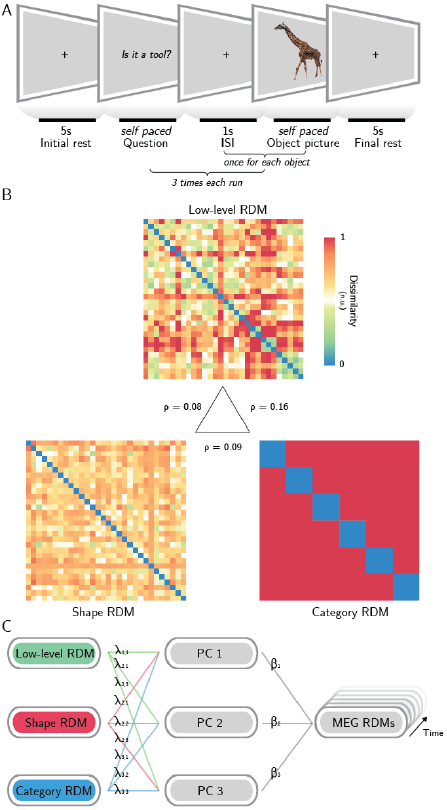
Methodological pipeline. A) Experimental design: subjects were asked to attend thirty object pictures during a semantic judgment task; B) representational dissimilarity matrices (RDMs) of three models (low-level features, shape and category) were employed to predict the MEG representational geometry - in the central triangle, Spearman correlation values between models are reported. With Relative Weights Analysis (C), MEG RDMs were predicted using three orthogonal principal components (PCs 1-3) obtained from the models, and the resulting regression weights were back-transformed to determine the relative impact of each model on the overall prediction when controlling for the impact of model collinearity (see Methods). Photo taken and edited from http://pixabay.com, released under Creative Commons CC0 license.

Thus, we first assessed the collinearity between the three models, expressed as the Spearman correlation between the model RDMs (Figure 2B). The low-level and categorical models have a correlation of ρ = 0.16, the shape model has ρ = 0.09 correlation with the categorical model, and ρ = 0.08 correlation with the low-level one.

Then, RWA (a method that controls for model multicollinearity in multiple regression – see Methods) was performed within a sensor space searchlight, resulting for each subject in three maps that report the time courses of the metric ε for each sensor, i.e., the proportional contribution of each model across time. Single-subject maps were then aggregated in group-level z-maps for each model (see Methods), corrected for multiple comparisons and divided in 50ms-long time bins for displaying purposes. Only the sensors whose corrected z-values were significant in the entire bin were retained, as displayed in Figure 3 (black dots mark significant sensors).

**Figure 3.**
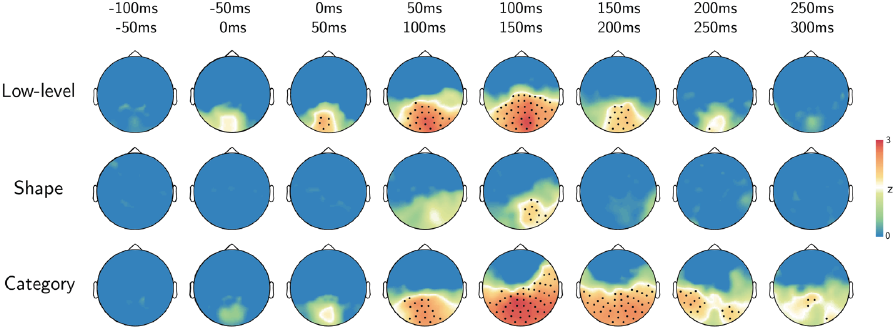
Results. Topographic plots of the group-level z-maps. Top-row reports the time bin. Black dots stand for significant channels within all the time-bin (p < 0.05, TFCE corrected).

Results show that the model based on low-level features (contrast and spatial frequencies) is significant at early stages after stimulus presentation (0-50ms) in a cluster of posterior and medial sensors. This cluster expands in the lateral and anterior directions, reaching a maximum in the 100-150ms interval, when most of the posterior sensors are significant. Shape features are instead restricted to right posterior location in the 100-150ms interval, and do not reach significance in the remainder of sensors and time bins. The category-based model is significant in medial and posterior sensors starting at 50-100ms. The cluster expands to most of the posterior and lateral sensors, with a maximum spatial extent between 100 and 200ms, then restricting to the posterior and lateral sensors in the 200-250ms time bin. A cluster of right posterior sensors shows significant weights for the three models in the 100-150ms time bin only. None of the models was significant in the remaining parts of the time course (before stimulus onset and after 300ms).

## Discussion

The visual machinery is a general-purpose system, relying on different representations that often are collinear or interact to each other. Here, by taking into account model collinearity, the spatiotemporal dynamics of joint feature processing within the human visual system were revealed to assess the relative contribution of low-level, shape and category features in predicting MEG-based representations. We observed both a temporal and spatial co-occurrence of low-level, shape and categorical processing, early in time (100-150ms) in posterior sensors. Specifically, we showed that a) low-level features (i.e., contrast and spatial frequencies) are processed early (0-50ms) after stimulus onset within posterior MEG sensors, spreading in time from medial to lateral locations; b) shape coding is limited within a few right posterior sensors in a brief time window (100-150ms) and co-occurs with low-level and categorical processing; c) categorical representation emerges later than the onset of low-level processing and is more prolonged, but spreads within a similar pattern of sensors.

Our results demonstrated that in the 100-150ms interval after stimulus onset, these features are processed concurrently, thus suggesting that object discrimination may result from independent parallel processing (i.e., orthogonal feature-based descriptions processed with similar temporal dynamics), rather than from a strict feed-forward hierarchy. The observed spatiotemporal overlap is in line with previous neuroimaging evidence showing that category and shape are processed within the same visual regions^14,19^, and can be decoded in the 130-200ms time window within the high-level visual cortex, as shown in a combined fMRI-MEG study, which just focused on body parts and clothes ^20^. Here we employed a model-based approach which also embedded low-level features, and sampled stimuli from a broader set of categorical classes. In addition, we introduced RWA to overcome multicollinearity, which was not explicitly addressed in previous studies.

Of note, our results raise questions concerning the role of shape in categorization. The synchronization between the three models in our data occurs in a time window (100-150ms) that overlaps with those of perceptual organization (70-130ms) and categorical recognition of visual information (>130ms), as indicated by previous neurophysiological and functional studies in both human and nonhuman primates^21,26^.

Whether shape processing is needed to recognize and classify objects in a scene has not been clarified yet. The classical view that considered shape essential to recognition^27^ has, however, being challenged by the success of several appearance-based computational models that could perform object recognition by relying on low-level features only ^28^. Since object segmentation occurs during passive natural image viewing^29^ and controls scene reconstruction^23^, shape analysis can be similarly triggered by object viewing also in a task for which shape is not explicitly relevant. Thus, our observation has at least two possible explanations: a) shape processing is to some extent necessary for categorization or, alternatively, b) it is not, but it is an automatic process occurring even when not overtly required by the task. The former hypothesis may, however, not be consistent with our results that show categorical representations occurring earlier than shape-based representations. In addition, the latter case would be in line with evidence suggesting that the extraction of object affordances – i.e., shape-related features which are able to facilitate or even trigger actions – is a fast and automatic process^30,31^. However, a conclusion on this topic can be reached only by further studies involving task modulation ^32^. Of note,task is able to influence the strength of object processing late in time (>150ms: ^33^).

Another interesting result is the early emergence (from 50ms) of categorical processing within the same pattern of sensors that also encode contrast and spatial frequencies. As mentioned before, object recognition has been described as occurring at 150ms or later^26^. We observed category representations within posterior sensors well before (even accounting for the temporal smoothing potentially introduced by the searchlight procedure). Early occurrence of categorical processing has been observed also in previous MEG studies^15,33^.

In the past years, mounting evidence revealed a top-down control of neurons in the early visual cortex^22,34-38^. Moreover, in a series of elegant studies^23,39,40^, Neri found psychophysical evidence of a top-down predictive mechanism, comprising a progressive refinement of local image reconstruction driven by global saliency or semantic content. At the macroscale, the effects of this mechanism imply that both local (i.e., low-level features) and global (i.e., object-related) representations should be retrieved early in time (<150ms) within the visual cortex. Our results, which show early (from 50 to 200ms), overlapping patterns for low-level and categorical processing in posterior MEG sensors, are in line with this view. However, further research is needed to directly test the causal role of top-down feedbacks in controlling low-level processing within the occipital cortex, which falls beyond the original scope of this work.

A further general remark should be made. As mentioned before, multicollinearity is a pervasive property of our surrounding environment. Indeed, one of the most fascinating features of our visual system is the way it deals with correlated statistics within the natural domain, to optimally represent the retinal input^41^, and to make sense of the external world, through the mean of learning and generalization. Indeed, visual correspondences are the mechanism we used to evolve more abstract, categorical representations^42^. However, from the researcher perspective, this leads to an extreme effort in balancing dimensions of interest, or in developing orthogonal models. In addition, two further aspects should be considered: first, as shown empirically ^12^, since different stimuli typically vary within multiple dimensions simultaneously, it is almost impossible to isolate a single dimension of interest; second, the effort in building orthogonal competing descriptions increases with the number of tested models.

Several methods have been proposed to overcome models collinearity issues when studying brain activity (for a review, see: ^43^). Within the field of neuroimaging, Lescroart, et al. ^44^ employed a variance partitioning approach (the same method, in the domain of multiple linear regression, is known as commonality analysis – as also employed in the MEG field^33^), which aims at determining the explained variance for any possible subset of the models. While this analysis is able to estimate the variance unique to each partition, its main drawback is that partitions grow exponentially with the number of models: since there are 2^p^ — 1 subsets for *p* predictors, just exploring the impact of 5 models generates 31 different subsets. In light of this, even comparing a low number of models would end up in a computationally intensive process and in the challenging task of interpreting and discussing a huge number of sub-models. Moreover, the partitions related to variance shared by different models can occasionally be negative, and the interpretation of these negative components is still matter of debate^45^. From this perspective, RWA is an attractive alternative, as it estimates the relative, non-negative weight of each model and does not imply to discuss more models or components than those initially considered.

Indeed, relative weights reflect in a suitable manner the proportional impact of each variable on the prediction of brain activity and - if the predictors are standardized - sum up to the total explained variance^16^. However, some limitations also affect RWA: the most relevant is that estimated weights are not invariant to the orthogonalization procedure employed. Though, it has been proven that, the more the orthogonal variables approximate the original variables, the more reliable the estimated weights become (for a deeper treatment of the topic, see: ^16^). Therefore, RWA may represent a fast and appealing recipe to deal with model multicollinearity within the neuroimaging field, especially when three or more models are compared.

In conclusion, this study reveals the spatiotemporal dynamics of object processing from a model-based perspective, providing evidence in favor of an integrated perceptual mechanism in object representation.

## Methods

### Participants

Sixteen healthy right-handed volunteers (5F, age 27 ± 2) with normal or corrected to normal visual acuity participated in the study. All subjects gave informed consent to the experimental procedures and received a monetary reward for their participation. The study was approved by the Ethics Committee for research involving human participants at the University of Trento, and all the experimental procedures were conducted in accordance with the Declaration of Helsinki.

### Stimuli

Visual stimuli were color pictures representing thirty different objects from six semantic categories (fruits, vegetables, animals, birds, tools, vehicles). Stimuli were presented using MATLAB and the Psychophysics Toolbox^46^, and were projected on a translucent screen placed at about 130cm from the participant, using a Propixx DLP projector (VPixx technologies), with a refresh rate of 60 Hz and a resolution of 1280×1024 pixels.

### Task and design

The experiment was organized in eight runs, each consisting of three blocks (see Figure 2A). In each block, the thirty images were presented in randomized order, and participants were engaged in a semantic judgment task to ensure that they focused the attention on the stimuli^47^. At the beginning of each block, a binary target question (e.g., “Is it a tool?”) was shown; once subjects read the questions, they prompted the start of the block by pressing a button on a keyboard. Within each block, subjects answered (yes/no) to the question presented at the beginning using the keyboard. All pictures were presented 24 times, with a different target question for each repetition. 5s-long resting periods preceded and followed each block, and 1s-long resting periods followed the behavioral response to each stimulus within a block. During the resting periods, subjects had to fixate a black cross, displayed in the center of the screen. The order of the questions was randomized across participants.

### Models

In order to predict MEG representational geometries, three different descriptions were built, representing different physiologically relevant properties of the objects seen by the subjects (see Figure 2B). First, a low-level model, which captures the arrangement of spatial frequencies in a V1-like fashion, was employed: a GIST^28^ descriptor for each stimulus was derived by sampling (in a 4×4 grid) the responses to a bank of isotropic Gabor filters (8 orientations and 4 scales). The descriptor (consisting of a vector with 512 elements) of each stimulus was then normalized and compared to each other stimulus using the pairwise correlation distance (1 - Pearson’s r). Second, a shape model was computed.Similarly to previous neuroimaging investigations on the same topic^48,49^, the medial-axis transform^18^ was extracted from each manually segmented and binarized object silhouette. Then, shock-graphs skeletal representations were built, and their pairwise dissimilarity was computed using the ShapeMatcher algorithm (http://www.cs.toronto.edu/͂dmac/ShapeMatcher/; Van Eede, et al. ^50^), which estimates the minimum deformation needed in order to match two different shapes^51^. Finally, the thirty stimuli were described based on their semantic category, obtaining a binary categorical model.

### MEG data acquisition

MEG data were recorded using an Elekta VectorView system with 306-channels, 204 first order planar gradiometers and 102 magnetometers (Elekta-Neuromag Ltd., Helsinki, Finland), located in a magnetically shielded room (AK3B, Vakuumschmelze, Hanau, Germany). The sampling rate was 1kHz. Head shapes were recorded from each participant immediately before the experiment, using a Polhemus Fastrak digitizer (Polhemus, Vermont, USA) recording the position of fiducial points (nasion, pre-auricular points) and around 500 additional points on the scalp. MEG data were synchronized with experiments timing by sending four different triggers at question presentation, first button press (after question), stimulus presentation and stimulus-related behavioral responses (button presses), respectively.

### MEG data preprocessing

MEG data preprocessing was performed using the Fieldtrip toolbox^52^. First, a bandpass (1-80 Hz) and a notch (50 Hz) 4^th^ order Butterworth IIR filters were applied to the data^53^. Filtered signals were then cut in epochs from 500ms before to 1s after stimulus onset and resampled at 400 Hz. Subsequently, data were visually inspected according to a set of summary statistics (range, variance, maximum absolute amplitude, maximum z-value) to search for trials and channels affected by artifacts, using the procedure for visual artifact identification implemented in Fieldtrip; trials marked as bad were rejected and noisy sensors were reconstructed by interpolating their spatial neighbors. On average, 8% of the trials and 10% of the channels were rejected for each subject.

### Searchlight analysis

A searchlight analysis was performed using CoSMoMVPA^54^, retaining the MEG data from the gradiometers only. First, the time-locked patterns for the individual trials were reduced to thirty pseudotrials (one for each stimulus)^55^. Searchlights were then defined for each time point of the pseudo-trials using a spatial and temporal neighboring structure^56^. Each searchlight included 10 dipoles (pairs of combined gradiometers) in the spatial domain, and each time point plus the ten preceding and following it (i.e., 21 time points, 52.5ms) in the temporal domain. Within each spatiotemporal searchlight, a time-varying representational dissimilarity matrix (RDM) was derived for the MEG data by computing the pairwise correlation distances between pattern of responses to the thirty stimuli^57^; prior to computing the RDM, stimulus-specific activity patterns were normalized (z-scored).

### Relative Weights Analysis (RWA)

In order to estimate how well each model RDM was related to MEG representational geometries, a multiple linear regression for each subject and each spatiotemporal searchlight was performed. Since some of the three models RDMs are significantly correlated (see Results) the Relative Weights Analysis (RWA), introduced by Johnson ^16^, was adopted. This method does not identify the impact of each model to the prediction of a dependent variable in isolation, as in common multiple linear regressions, but considers also how each model relates to (i.e., is correlated with) the others. The metric on which RWA relies is called epsilon (ε) and reflects both the unique contribution of each model and its impact when all the other models are considered.

The RWA procedure is graphically synthetized in Figure 2C. Basically, the models RDMs were first orthogonalized, by performing a Principal Component Analysis (PCA), and the RDMs from each spatiotemporal searchlight were regressed on the so obtained orthogonal versions of the models RDMs. Then, the regression coefficients were related back to the original model RDMs by regressing the orthogonal RDMs also on the models RDMs. Finally, for the *j-th* model, epsilon was calculated as:

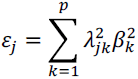

where *p* is the number of models, 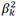 is the variance (i.e., the squared standardized regression coefficient) in each searchlight RDM accounted for by the *k-th* orthogonal RDM, and 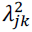 is the variance in the *j-th* model accounted for by the *k-th* orthogonal RDM.

### Statistical analyses

The RWA analysis, performed within the spatiotemporal searchlights as described above, provided a time course of the metric (ε) for each sensor and time point. To estimate the group-level spatiotemporal distribution of weights for each of the three models, a one sample non-parametric test was performed, using a null distribution generated with 100,000 permutations (rank test), as implemented in CoSMoMVPA. Correction for multiple comparisons was made at cluster-level using a threshold-free method (TFCE: ^58,59^). Z-values corresponding to a corrected p-value of 0.05 (one-tailed) were considered significant.

## Author contributions

P.Papale, A.L. and E.R. conceived the study; E.R., G.H, A.R. and A.L. designed the experimental paradigm; A.L., A.R., L.T., G.M. performed the experiments, P.Papale., A.L., and M.B. analyzed the data; P.Papale and A.L. wrote the manuscript; E.R, L.C., G.H., A.R., L.T. and P.Pietrini critically revised the manuscript.

## Additional information

The authors declare no competing financial and non-financial interests.

